# *Pseudomonas syringae* socially-induced swimming motility requires the molybdenum cofactor

**DOI:** 10.1101/2024.05.20.595025

**Authors:** Zichu Yang, Bryan Swingle

## Abstract

Social interactions among bacteria can induce behaviors that affect their fitness and influence how complex communities assemble. Here we report a new socially-induced motility behavior that we refer to as baited expansion in *Pseudomonas syringae* pv. tomato DC3000 (*Pst* DC3000), a plant pathogenic bacterium. We found *Pst* DC3000 displayed strongly-induced swimming motility towards nearby colonies of *Dickeya dianthicola* or *Escherichia coli*. We developed a controlled system to visualize and characterize the development of baited expansion. Our results provide evidence that baited expansion behavior occurs in response to a chemical gradient established and maintained by the bait colony. We also found this behavior correlated with distinct transcriptional profiles and identified molybdenum cofactor as a crucial factor in facilitating the baited expansion behavior.

## Introduction

Soil bacteria, epiphytes, endophytes, and pathogens have the potential to encounter many other microbes in nature. Bacteria evolved to perceive and communicate with related (i.e., kin) and unrelated individuals (non-kin) and to modify their behavior in response (Waters and Bassler 2005; Bodman et al. 2008; Humphries et al. 2017; Liu et al. 2017). Several bacterial species also show the capacity for kin discrimination in their social behaviors, these depend on genetic relatedness between individuals and often help to direct benefits to kin (Senior 1977; Cornforth and Foster 2013; Stefanic et al. 2015). Diene’s line observed in *Proteus mirabilis* (Senior 1977; Gibbs et al. 2008), contact dependent inhibition (Aoki et al. 2005, 2010; Hayes et al. 2014) and similar boundary-forming observed in swarming *Bacillus subtilis* (Stefanic et al. 2015) are all types of social behaviors where the differential treatment towards kin and non-kin can be visually observed on typical semi-solid growth media.

*Pseudomonas syringae* pv. *tomato* DC3000 (*Pst* DC3000) is a model plant pathogenic bacterium that infects and causes disease in tomato (*Solanum lycopersicum*) and *Arabidopsis thaliana* (Buell et al. 2003). *Pst* DC3000 and other *P. syringae* strains, are globally dispersed, cosmopolitan bacteria that can be found co-occurring with a wide-range of bacteria in many types of environments (Morris et al. 2008). In this study, we used two genera from the family Enterobacteriaceae as models to stimulate *Pst* DC3000 socially-induced motility. *Pst* DC3000 has potential crosstalk with the quorum sensing systems of enterobacteria (Hosni et al. 2011) and also possesses a type VI secretion system (T6SS) that contributes to survival in inter-species competition against entrobacteria and yeast (Haapalainen et al. 2012). We previously found *Pst* DC3000 exhibited strongly induced expansion when cultured on swimming media with a second colony of unrelated bacteria such as *Xanthomonas campestris*, *Erwinia amylovora*, *Dickeya dianthicola* and *Escherichia coli* (Reeve et al. manuscript submitted). In contrast, *Pst* DC3000 had significantly less expansion in the direction of a second *Pst* DC3000 colony and grows radially without other colonies present (Reeve et al. manuscript submitted). Examples of socially-induced motility have been reported in other bacteria including *Pseudomonas aeruginosa*, *Streptomyces venezuelae* and *Bacillus subtilis* with diverse triggering mechanisms and interacting partners (Jones et al. 2017; Liu et al. 2018; Limoli et al. 2019; Shepherdson and Elliot 2022; Yarrington et al. 2024). Chemotaxis, antibiotic stress, and iron stress have been identified as important factors in development of these colony expansion phenotypes (Liu et al. 2018; Shepherdson and Elliot 2022; Yarrington et al. 2024).

In this study we report a new socially-induced swimming behavior, which we can induce and visualize by culturing *Pst* DC3000 in a soft agar medium inoculated with a different species of bacteria at a second location. We show this behavior does not occur between clonal *Pst* DC3000 swimming colonies and results from motility regulation rather than modulating growth rate. We found that subpopulations showing the induced motility have distinct transcriptional signatures and we used this information to identify a gene involved with molybdenum cofactor biosynthesis that is necessary for proper execution of the induced motility. We confirmed that molybdenum cofactor biosynthesis is necessary for the development of this behavior by targeting another gene that contributes to molybdenum cofactor biosynthesis and found induced social motility was also disabled with this mutant.

## Results

### Two enterobacteria species induce *Pst* DC3000 swimming colony expansion and directional bias

*Pst* DC3000 use 1-5 polar flagella to power swimming motility, which enables them to expand radially from a central inoculation point on low-percentage agar swimming medium (Sampedro et al. 2015) and form symmetrical colonies when they are grown in isolation. However, we found that *Pst* DC3000 colony expansion increased when cultured on swimming media with a second clone of unrelated bacteria such as *E. coli* and *D. dianthicola*. We developed a controlled swimming assay to study this behavior, in which wild-type *Pst* DC3000 was inoculated 5 mm from a non-motile strain of *E. coli* and *D. dianthicola* to act as bait to induce *Pst* DC3000 social motility (Fig. 1A and B). Baited expansion occurs in response to motile bait strains, but we used non-motile bait to compensate for intrinsic growth and motility differences between the bait species, which simplified comparison of *Pst* DC3000 responses. Using this assay, we saw *Pst* DC3000 display similar colony shapes in response to *E. coli* and *D. dianthicola,* which were markedly different from those that formed in response to non-motile *Pst* DC3000 bait or the unbaited control (Fig. 1C). When baited by either of these enterobacteria species, the *Pst* DC3000 colonies occupied a larger area and were less turbid on the periphery, with the edge most proximal to the bait becoming progressively less turbid to the point of being nearly indiscernible (Fig. 1A and B). An additional line of dense cells also appeared on the portion of *Pst* DC3000 colony proximal to the bait, which was absent when the colony was grown in isolation or when baited by another *Pst* DC3000 colony (Fig. 1A and B).

**Figure 1.**
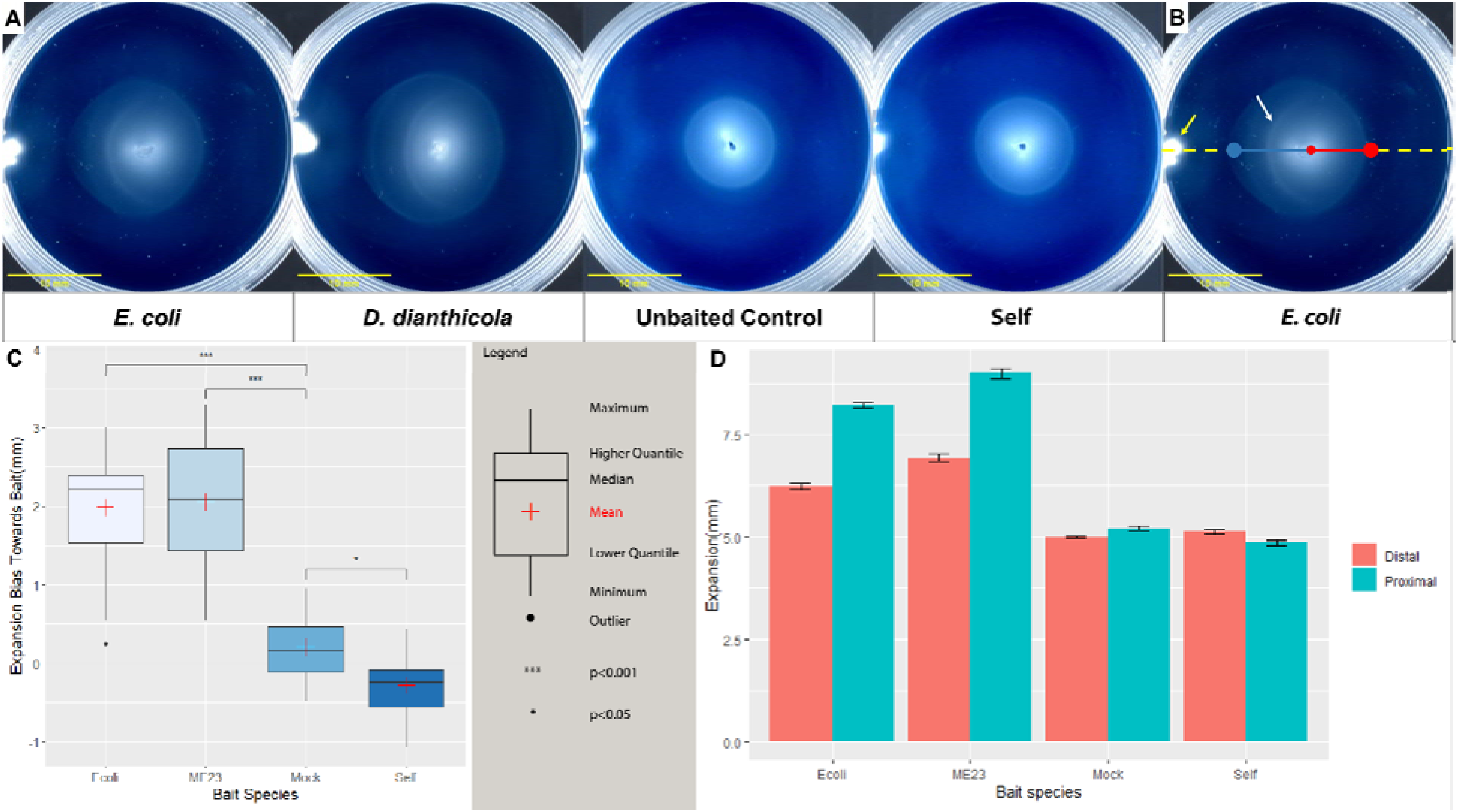
Baited expansion behavior of *Pst* DC3000 swimming colonies. (A) Representative results of baited swimming assays at 48 hpi for *Pst* DC3000 (PS1) paired with indicated baits. Bait species from lef to right are *E. coli* (BMS533), *D. dianthicola* (ME23 derivative, PS1255), Unbaited control, *Pst* DC3000 (Self, PS398); Images were processed with ImageJ enhance contrast function. (B) Graphical explanation of method used to quantify *Pst* DC3000 swimming colony expansion characteristics. Small red dot in center shows the location of *Pst* DC3000 colony inoculation point, which is 15 mm from location of bait (yellow arrow). Proximal and distal expansion distances were measured along the dashed line going through *Pst* DC3000 and bait inoculation points. Solid blue and red lines represent the length of the proximal and distal expansion distances, respectively. Large blue and red dots show the proximal and distal colony edges, respectively. White arrow points to the dense line of cells appearing in enterobacteria baited *Pst* DC3000 colonies. (C) Average *Pst* DC3000 swimming colony expansion distances on proximal and distal sides in response to the indicated bait species. Error bars show the standard error. Proximal expansion is significantly greater compared to distal expansion when baited by *E. coli* and *D. dianthicola* (p < 0.001, paired t-test, N=27). (D) Distribution of measured expansion bias in all 4 groups of baited swimming assays (student’s t-test, N = 27).

**Table 1.** Strains and plasmids used.

| Strain or plasmid | Relevant characteristic(s) | Reference |
| --- | --- | --- |
| Strains |  |  |
| PS1 | *P. syringae* pv. *tomato* DC3000 wild type | Buell et al., 2003 |
| PS129 | *P. syringae* pv. *tomato* DC3000 wild type with pBS60 control plasmid | Markel et al., 2016 |
| PS398 | *P. syringae* pv. *tomato* DC3000 Δ*cheA2* | Clarke et al. 2016 |
| BMS378 | *D. dianthicola* ME23 | Ma et al, 2019 |
| PS1255 | *D. dianthicola* ME23 Δ*cheA* MotB.p.Y317X | This work |
| BMS533 | *E. coli* JW1877 *cheA::kan* Keio collection CGSC 9563 | Baba et al., 2006 |
| PS1439 | *E. coli* JW1877 *cheA::kan* with pBS60 | This work |
| PS1854 | *P. syringae* pv. *tomato* DC3000 ΔPSPTO_5639 | This work |
| PS1855 | *P. syringae* pv. *tomato* DC3000 Δ*moaA* | This work |
| PS1856 | *P. syringae* pv. *tomato* DC3000 ΔPSPTO_5639 with pIY54 | This work |
| PS1857 | *P. syringae* pv. *tomato* DC3000 ΔPSPTO_5639 with pBS60 | This work |
| PS1858 | *P. syringae* pv. *tomato* DC3000 Δ*moaA* with pIY52 | This work |
| PS1859 | *P. syringae* pv. *tomato* DC3000 Δ*moaA* with | This work |
|  | pBS60 |  |
| PS1899 | *P. syringae* pv. *tomato* DC3000 $\Delta moeA$ | This work |
| PS1900 | *P. syringae* pv. *tomato* DC3000 $\Delta moeA$ with pIY56 | This work |
| PS1901 | *P. syringae* pv. *tomato* DC3000 $\Delta moeA$ with pBS60 | This work |
| PS909 | *P. syringae* pv. *tomato* DC3000 with Tn7 integrated GFP | This work |
| **Plasmids** |  |  |
| pBS46 | pBBR1MCS5 derivative, Gateway compatible broad-host range expression vector with nptII promoter, Gm<sup>r</sup> | Wei et al., 2007 |
| pBS60 | P<sub>nptII</sub>::empty control vector for pBS46 based expression, Gm<sup>r</sup> | Swingle et al., 2008 |
| pK18mobsacB | Allelic exchange suicide vector; lacZ mob sacB; Km<sup>r</sup> Suc<sup>s</sup> | Schafer et al., 1994 |
| pR6KT2GW | Allelic exchange suicide vector, Gateway<sup>TM</sup> BP clonase compatible, gus<sup>+</sup>, SacB<sup>+</sup>, Gm<sup>r</sup>, Cm<sup>r</sup> | Stice et al., 2020 |
| pIY11 | pR6KT2GW based *cheA* deletion construct (fused with XhoI-PstI) | This work |
| pIY51 | pK18MobSacB based *moaA* deletion construct, | This work |
|  | Km<sup>r</sup> |  |
| pIY52 | pBS46 based nptII::*moaA* expression construct,<br>Gm<sup>r</sup> | This work |
| pIY53 | pK18MobSacB based PSPTO_5639 deletion<br>construct, Km<sup>r</sup> | This work |
| pIY54 | pBS46 based nptII::PSPTO_5639 expression<br>construct, Gm<sup>r</sup> | This work |
| pIY55 | pK18MobSacB based *moeA* deletion construct,<br>Km<sup>r</sup> | This work |
| pIY56 | pBS46 based nptII::*moeA* expression construct,<br>Gm<sup>r</sup> | This work |

We quantified the baited expansion phenotype by determining the directional bias and the expansion induction in these swimming assays. The directional bias was calculated as the difference between the proximal expansion distance relative to the distal expansion distance (Fig. 1B and C). The expansion distance describes the distance between the inoculation point and the proximal and distal edges of *Pst* DC3000 colonies (Fig. 1D). Enterobacterial baits induced *Pst* DC3000 colony expansion while the *Pst* DC3000 bait did not (p < 0.001, paired student’s t-test, N = 27) (Fig. 1A, C and D). At 48 hours post inoculation (hpi), *Pst* DC3000 colonies baited by enterobacteria exhibited 8 to 9 mm proximal expansion and 6 to 7mm distal expansion, while both sides averaged to around 5 mm when baited by either non-motile *Pst* DC3000 or the unbaited control. *Pst* DC3000 colonies baited by both enterobacteria strains exhibited ten times the directional bias of the unbaited control (Fig 1C), with proximal side of the colony expanding about 30% more than the distal side (Fig. 1D). *Pst* DC3000 colonies baited by non-motile *Pst* DC3000 exhibited a negative directional bias (Fig. 1C), which is consistent with our previous finding that *Pst* DC3000 colonies repel expansion (Reeves et al., manuscript submitted).

### The magnitude of *Pst* DC3000 baited expansion correlates with exposure to bait colony derived stimulus

The directional bias we observed with the baited *Pst* DC3000 swimming colonies seemed to occur because of increased expansion in parts of the colonies closer to the bait. This observation suggested that the baited *Pst* DC3000 cells were responding to a gradient established by the bait colony with cells closer to the bait reacting to more stimulus as in a dosage-dependent response. If this is true, then giving the bait more time to establish this gradient should increase the *Pst* DC3000 response overall. To test this, we compared the magnitude of baited expansion exhibited by *Pst* DC3000 when the *E. coli* bait and *Pst* DC3000 were inoculated simultaneously to that when the bait colony had been allowed to grow in the assay medium for 24 hours prior to inoculating the *Pst* DC3000 colony. *Pst* DC3000 swimming colonies baited by a pre-inoculated *E. coli* colony showed increased expansion induction characteristics compared to that triggered by an *E. coli* colony inoculated at the same time as *Pst* DC3000, including wider colonies (i.e. more lateral expansion of the bait proximal portion of the colony) and enhanced dense line formation in the parts of the colonies most proximal to the bait (Fig. 2A).

**Figure 2.**
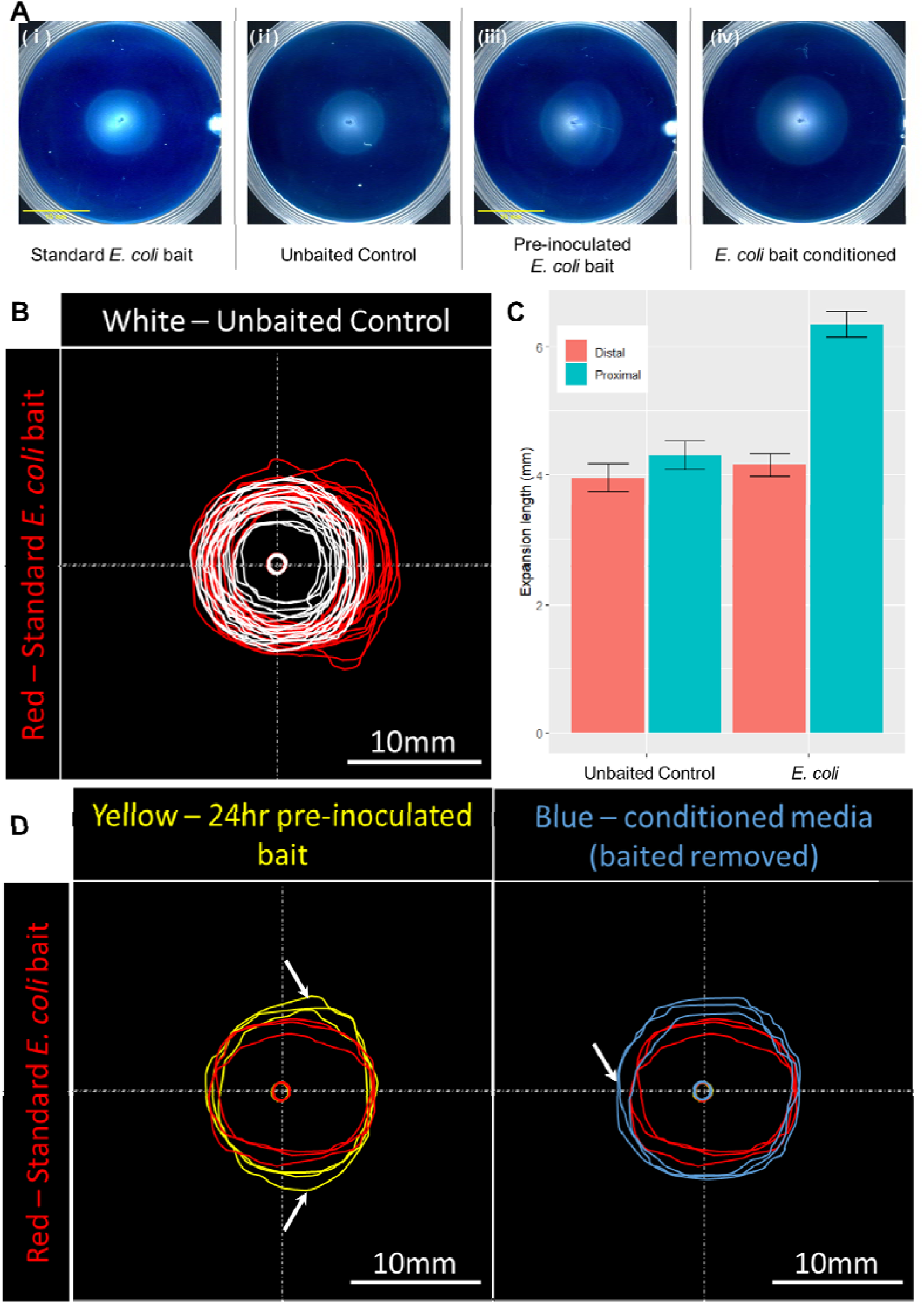
Baited expansion behavior of *Pst* DC3000 colonies in *E. coli* bait conditioned swimming medium. (A) Representative colony morphology of *Pst* DC3000 (PS129) at 48 hpi in baited swimming assays with following specific conditions: (i) standard baited swimming assay with *E. coli* bait (PS1439) inoculated at the same time as the *Pst* DC3000 colony (N = 15); (ii) unbaited control (N = 13) (iii) *E. coli* bait (PS1439) inoculated 24hr before the *Pst* DC3000 colony (N =3); (iv) baited swimming assay with swimming medium conditioned by *E. coli* (PS1439) growth on nitrocellulose membrane for 72 hr, which was removed prior to *Pst* DC3000 inoculation (N =3). (B) Tracings of *Pst* DC3000 swimming colony shapes from (i) and (ii) (C) Quantification of proximal and distal expansion length of *Pst* DC3000 swimming colony at 48 hpi in gentamycin-supplied baited swimming assays (i) and (ii). Directional expansion bias was preserved despite addition of gentamycin. (D) Comparisons of 48 hpi *Pst* DC3000 swimming colony shapes across (i) (iii) and (iv). 24hr pre-inoculation of bait led to induced expansion in additional directions on the bait-proximal side of the swimming colony (white arrow, bottom-left panel). Bait conditioned swimming agar (bait-removed) induced colony expansion in *Pst* DC3000 symmetrically, resulting in colony shapes with extra expansion on the distal side (white arrow, bottom-right panel). In these tests we used *Pst* DC3000 transformed with pBS60 (Swingle et al. 2008a), so that we could use gentamycin resistance to select for *Pst* DC3000 to minimize contamination.

We manually recorded and overlayed swimming colony edges at 48 hpi to better compare the expansion shapes and areas of *Pst* DC3000 colonies exposed to various bait conditions (Fig. 2B and C). Overlay of colony edges confirmed that *Pst* DC3000 swimming colonies baited by a pre-inoculated *E. coli* colony showed increased expansion and increased directional bias (Fig. 2D).

We next tested whether baited expansion behavior requires the presence of a live bait colony or whether *Pst* DC3000 could respond to the stimulus without live bait present. To test this, we produced conditioned medium by growing an *E. coli* colony on a nitrocellulose membrane for 72 hours at the edge of the plate where the bait colony would normally have been inoculated, then removed the filter and recorded the size and shape of *Pst* DC3000 colonies 48 hours after they were inoculated in the conditioned medium. This allowed nutrients and metabolites to freely diffuse through the membrane while preventing bacteria from moving into the medium, and we confirmed there was no evidence of bait colony growth in the medium after the membrane was removed. We found that the resulting colonies expanded more than colonies growing in unconditioned swimming media, but without the directional bias observed with the bait colony present (Fig. 2D). This suggests that the conditioned medium contained sufficient quantities of bait stimulus to induce *Pst* DC3000 baited expansion behavior, but that the gradient had dissipated and *Pst* DC3000 could not track the original location of the *E. coli* colony (Fig. 2D). We also attempted to induce this behavior by adding supernatant of stationary phase *E. coli* liquid cultures to the swimming agar but could not detect any changes to *Pst* DC3000 swimming motility or colony expansion.

### *Pst* DC3000 baited expansion results from increased directional motility

*Pst* DC3000 form asymmetric colonies when baited by *E. coli*, with the proximal side occupying a larger area with an apparent lower cell density than the distal side of the colony (Fig. 2B). This asymmetric colony morphology can result from either altered growth, motility induction or a combination of both. To assess the role of growth in development of baited expansion behavior, we compared the number of viable cells in *E. coli*-baited *Pst* DC3000 colonies to unbaited controls. To estimate total number of cells we removed the colonized medium from the petri dishes, disintegrated the gel matrix using vigorous agitation with sterile glass beads, plated dilutions of this mixture, and counted colony forming units (CFU). Surprisingly, there was no detectable difference in the total number of cells regardless of bait presence (Fig. 3A, t-test, p > 0.1, N = 9). This suggests that baited *Pst* DC3000 does not receive an immediate fitness benefit under these conditions despite expanding more.

**Figure 3.**
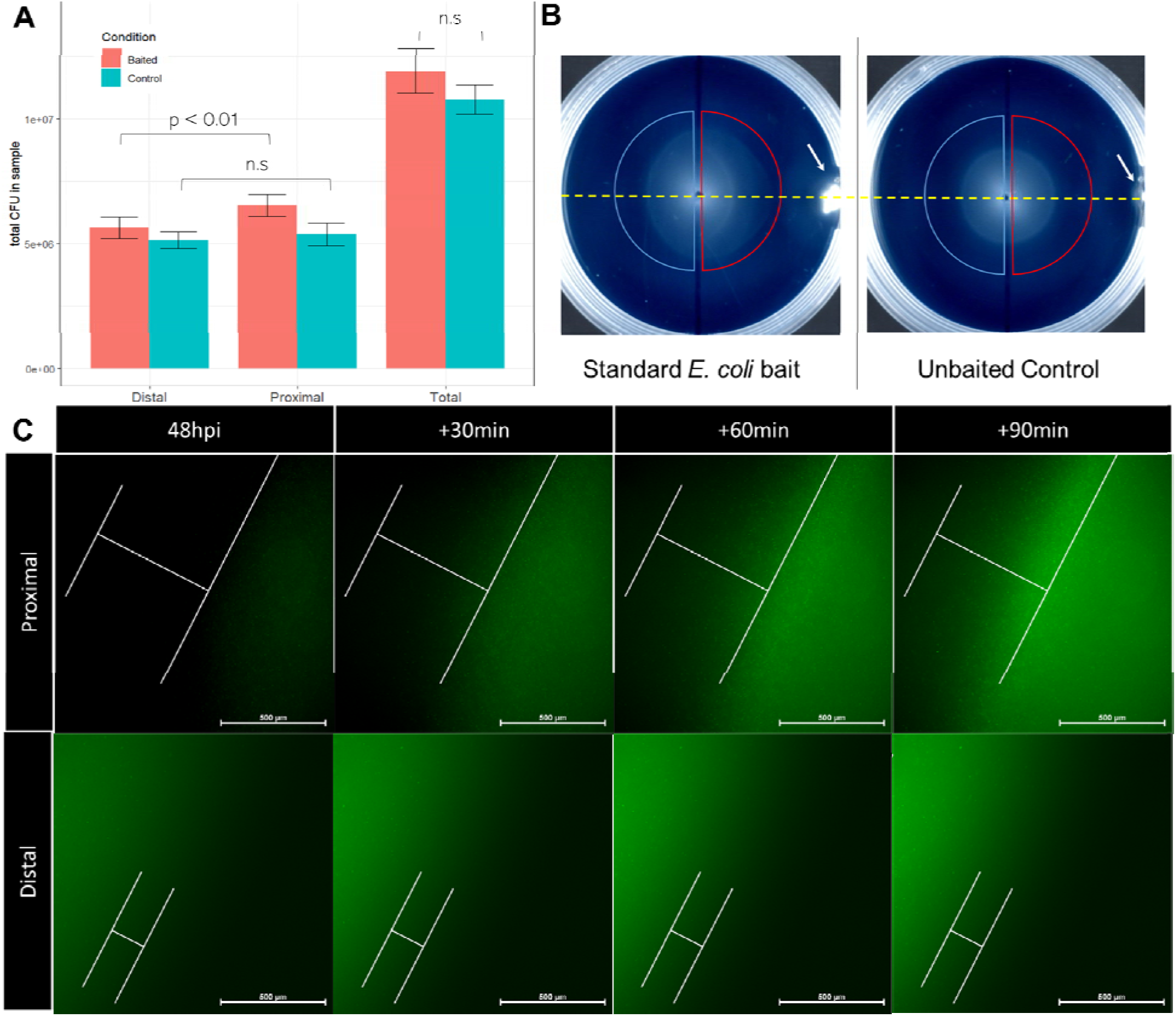
Analysis of growth and motility as factors contributing to baited expansion behavior. (A) Average number of CFUs from proximal and distal regions and combined total CFUs of baited and unbaited swimming colonies. Total CFUs were not significantly different between baited and control swimming colonies despite differences in colony area (p > 0.1, student’s t-test, N = 9). Proximal side of baited swimming colonies had more viable cells than distal side (p < 0.01, paired student’s t-test, N = 9) but there was no detectable difference in the number of viable cells in each side of unbaited control colonies (p > 0.1, paired student’s t-test, N = 9). (B) Graphical explanation of method used to determine the number of cells in proximal and distal regions of swimming colonies. Inoculation point for bait colony or sterile medium control are marked by white arrows. A line perpendicular to the dash line was marked on the bottom of each plate to separate the swimming colony into two regions: proximal to bait (red) and distal to bait (blue). (C) Time series images of the proximal and distal edges of GFP-tagged *Pst* DC3000 (PS909) swimming colonies at fixed coordinates visualized by epifluorescence microscopy beginning at 48 hpi. Approximate starting and final colony edges are marked by solid white lines. Proximal edge expanded 600 μm during the 90-minute incubation while distal edge moved less than 200 μm.

Next, we compared the number of *Pst* DC3000 cells in the proximal sides to the number in the distal sides of *E. coli*-baited *Pst* DC3000 colonies. To estimate total number of cells in each region we separated the distal and proximal sides of the baited colonies by cutting the colony on the line through the inoculation point and perpendicular to the axis of the directional bias, (Fig. 3B) and collected the cells from the medium as described above. We found that the proximal side of *E. coli* baited *Pst* DC3000 colonies contained 10% to 30% more viable cells than the distal side (Fig. 3A, paired t-test, p < 0.01, N = 9). These results suggest that there is a significant redistribution of cells within the colony during baited expansion, in which cells from proximal and distal parts of the colony move towards bait colony. Together with observed larger colony expansion, these data suggest that baited expansion is a product of induced motility rather than increased growth.

We then used fluorescence microscopy to examine the difference in expansion between the proximal and distal edges of *E. coli*-baited *Pst* DC3000 colonies. We recorded mass expansion (growth and motility) of *Pst* DC3000 cells expressing the green fluorescent protein (GFP) at the leading edges of *E. coli*-baited colonies over 90 minutes beginning at 48 hpi. We found the subpopulation at the proximal edge of the colony had higher cell density as well as overall more moving cells in comparison to the distal edge population (Video 1 and 2). In addition, we found the proximal edge of the baited swimming colony moved approximately 600 μm over 90 minutes while the distal edge of colony moved less than 200 μm during same time period (Fig. 3C).

### Sub-populations of baited *Pst* DC3000 swimming colonies are distinguishable at transcriptomic level

*Pseudomonas fluorescens*, a close relative to *P. syringae*, has specific transcriptomic responses depending on the species of neighboring colonies (Garbeva et al. 2011). Based on this, we hypothesized that proximity to bait could have similar impact on *Pst* DC3000 transcription and that comparison of transcriptomic responses among cells in different parts of baited colonies could be used to identify genes contributing to this socially affected behavior. We sampled sub-populations at the proximal and distal edges of *E. coli*- or *D. dianthicola*-baited *Pst* DC3000 swimming colonies (Fig. 4A), where the most extreme differences in expansion induction were observed, and assessed the transcriptome of these sub-populations by RNAseq. For comparison we also sampled the edge sub-populations from *Pst* DC3000-baited colonies and unbaited controls. Principal component analysis of differentially expressed genes among these sets of transcriptomes revealed that the proximal edge subpopulations in each enterobacteria-baited scenario displayed robustly unique expression patterns, as compared to the distal subpopulations, which are indistinguishable from each other or the edge population of an unbaited control colony (Fig. 4B). Out of 5891 total annotated genes in the *Pst* DC3000 genome, we found 270 differentially expressed genes (DEGs) between the proximal and distal subpopulations when baited by *E. coli* and 147 DEGs between these two subpopulations when baited by *D. dianthicola* (Supplemental Table S1). A total of 84 DEGs were shared between both bait conditions, and we found carbohydrate metabolic process as the strongest enriched GO terms among them (Fisher’s exact test, FDR p < 0.01).

**Figure 4.**
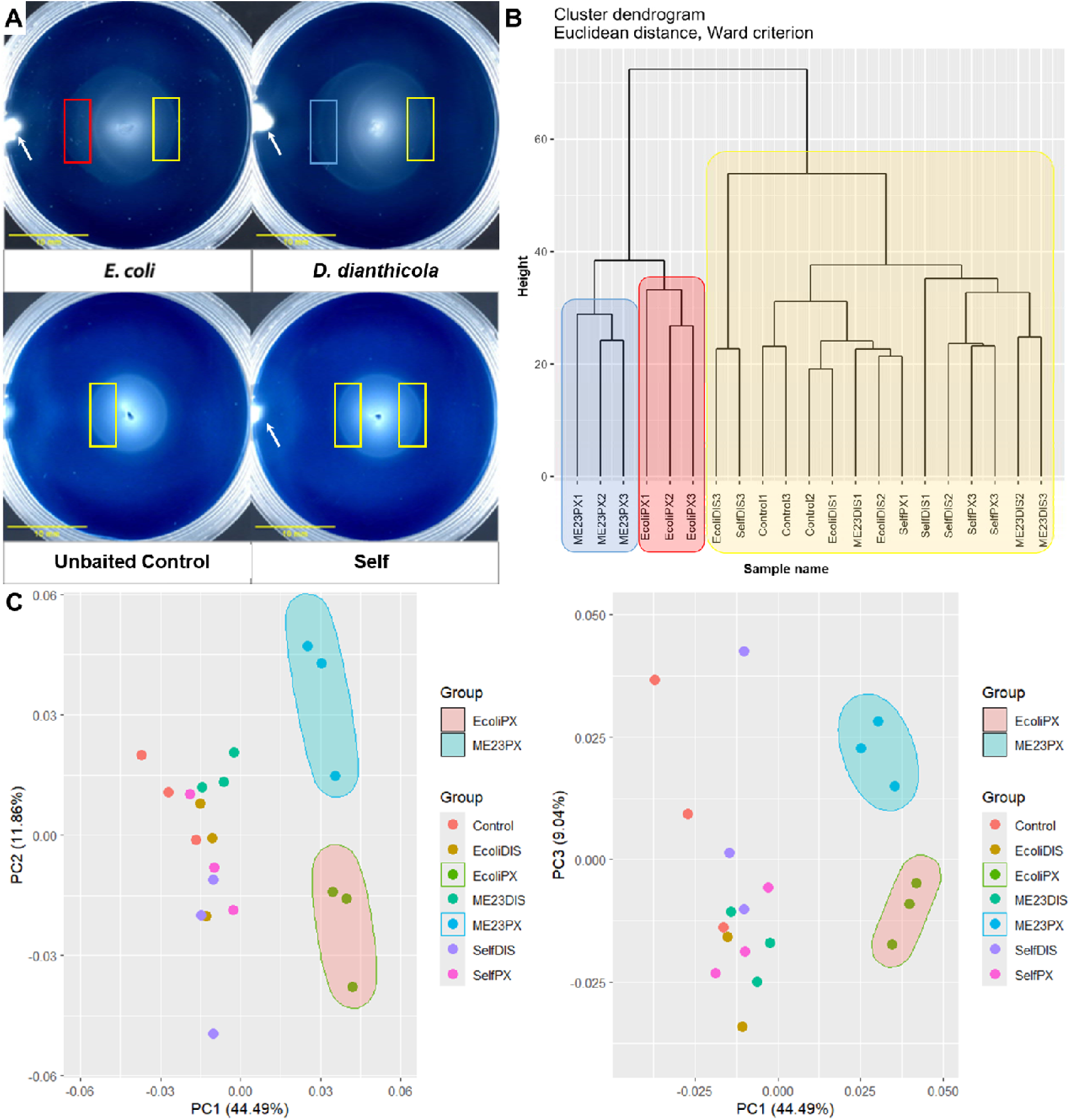
Edge sub-populations have distinct transcriptomic patterns. (A) Graphical explanation of sub-population sampling. White arrows indicate bait inoculation points. A gel extraction tool was used to sample 28 mm^2^ area (3.3 X 8.4 mm) at indicated parts of *Pst* DC3000 colonies 48 hpi. Color coding refers to cluster groups in panels B and C. (B) Clustering dendrogram of transcriptome of each sub-population of *Pst* DC3000 (PS1) swimming colonies. Samples from *E. coli* (BMS533) baited proximal edge (red) and *D. dianthicola* (PS1255) baited proximal edge (blue) are distinguishable from all other conditions (marked by yellow). (C) Principal component analysis of sample transcriptomes. Subpopulations proximal to enterobacteria bait are distinguished from subpopulations under other conditions by PC1 which accounts for 44.49% of difference in the transcriptomic patterns. PC2 and PC3 resolve transcriptional profiles of *E. coli* and *D. dianthicola* baited *Pst* DC3000.

*Pst* DC3000 swimming motility is predominantly regulated through control of flagellum expression and assembly. Flagellar activity and direction are controlled by the chemotaxis system (Sampedro et al. 2015) and cyclic di-GMP (c-di-GMP) (Pfeilmeier et al. 2016; Wang et al. 2019) but can also be modulated by many other environmental sensing systems like extacytoplasmic function sigma factor AlgU (Bao et al. 2020; Wang et al. 2020). *Pst* DC3000 has three annotated chemotaxis operons with the *che2* operon identified as the most essential for regulating swimming motility (Clarke et al. 2016). Surprisingly, we found only one gene related to cyclic di-GMP (PSPTO_0536), and one methyl-accepting chemotaxis gene (PSPTO_2448) in this shared DEG list (Supplemental Table. S1).

### Molybdenum cofactor biosynthesis genes are necessary for *Pst* DC3000 baited expansion

The RNAseq analysis showed that PSPTO_5639 (encoding hypothetical protein) had the strongest expression fold-change in the *E. coli* and *D. dianthicola* baited-proximal populations of *Pst* DC3000. We also found *moaA* (PSPTO_2516), which is 3 genes away from PSPTO_5639, had similar expression patterns. We confirmed the expression patterns of both genes by qRT-PCR (Fig. 5A). We used a genetic approach to test whether these genes have roles in motility or socially-induced baited expansion. We found no motility or behavioral differences between ΔPSPTO_5639 and wild type with the baited swimming assays. However, *moaA* deletion resulted in reduced swimming motility and a clear behavioral difference during baited swimming assay (Fig. 5B and C). At 48 hpi, expansion bias was reduced and the colony edge on the proximal side was well-defined in swimming *moaA* deletion strain compared to wild type. The wild-type phenotype was restored to the *moaA* deletion strain with *moaA* expressed from a plasmid (Fig. 5C and E).

**Figure 5.**
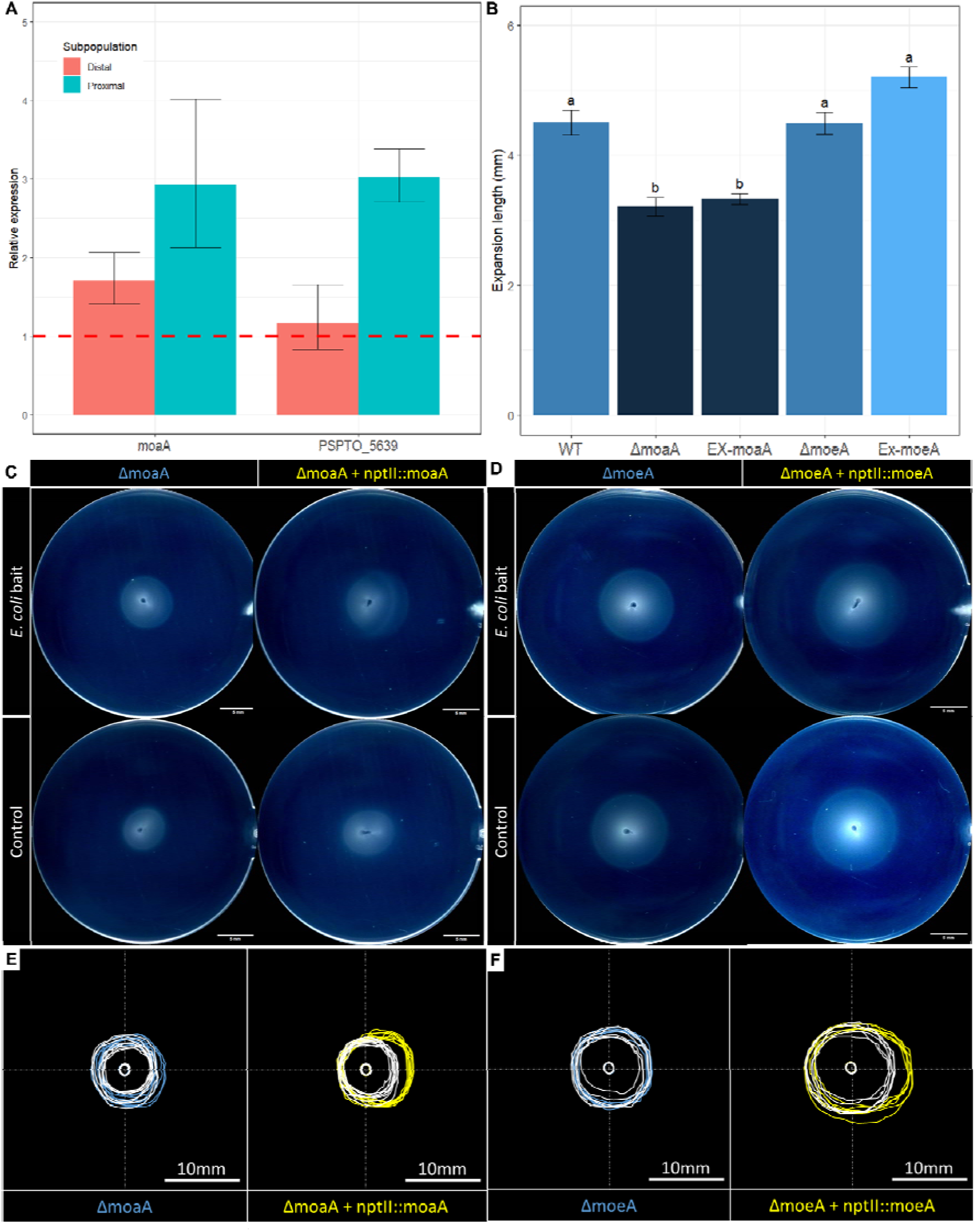
Molybdenum cofactor biosynthesis genes are required for baited expansion behavior. (A) qRT-PCR confirmation of context dependent differential gene expression of *moaA* (PSPTO_2516) and PSPTO_5639 in subpopulations of *E. coli* (PS1439) baited swimming *Pst* DC3000 (PS129) colonies. Red dash line represents relative expression level of *gyrA*, which was used to normalize expression of target genes. (B) Radius of unbaited swimming *Pst* DC3000, *moaA* and *moeA* deletion derivatives, and deletion derivates complemented with the indicated gene overexpressed from a plasmid. Letter groups were assigned by post hoc Tukey test. (C) Representative swimming colony shapes of *moaA* deletion strain with and without plasmid expressing *moaA*. (D) Representative swimming colony shapes of *moeA* deletion strain with and without plasmid expressing *moeA*. (E) Comparisons of 48hpi *Pst* DC3000 swimming colony shapes, 9 replicates for each condition. White line shapes are unbaited control colonies of the respective deletion strain. (F) Representative swimming colony shapes of *moeA* deletion strains. (E) Comparison of 48hpi *Pst* DC3000 swimming colony shapes, 6 replicates for each condition.

Annotation of *moaA* gene suggested that it functions as a GTP cyclase needed for biosynthesis of the molybdenum cofactor (Moco). This suggested that the changes in motility observed with the *moaA* deletion strain might be due to absence of Moco. To test this, we analyzed the swimming and socially-induced motility phenotypes of a strain with the *moeA* (PSPTO_2353) deleted, which is another gene necessary for Moco biosynthesis (Andreae et al. 2014; Leimkühler 2020) but was not regulated in response to baited or other conditions examined in our RNAseq experiment. Deletion of *moeA* resulted in loss of baited expansion behavior and eliminated the expansion bias, dense cell lines formation and the less-defined proximal colony edge in the baited swimming assay (Fig. 5D and F). All baited expansion behavior characteristics were restored by complementing the *moeA* deletion with *moeA* expressed from a plasmid (Fig. 5D and F). However, in contrast to the *moaA* deletion, the absence of *moeA* had no effect on unbaited swimming motility compared to wild type (Fig. 5B). Combined with the phenotypes of *moaA* deletion mutant, these data suggest Moco mediates the development of baited expansion behavior.

## Discussion

In this study we report a novel socially-induced motility behavior in *P. syringae*, which we are calling baited expansion. We characterized this behavior using its macroscopic colony-level qualities: induced expansion, directional expansion bias, formation of dense lines of cells within the colony and less defined colony boundary on the edge proximal to a second unrelated bait colony. We found this behavior occurs exclusively in interspecies interactions and the colony morphologies emerge primarily from induced motility rather than growth. Our investigation revealed that *Pst* DC3000 colony subpopulations have distinct transcriptomic profiles that depend on their location relative to the bait colony. We identified Moco biosynthesis genes as crucial contributors to the development of this behavior. Our data provide another example for the multicellularity of a motile bacteria colony where subpopulations are modifying their motility behavior based on social context.

We obtained three lines of evidence suggesting that *Ps*t DC3000 baited expansion behavior occurs in response to a chemical gradient established and maintained by the bait colony. First, *Pst* DC3000 colonies are asymmetric in the presence of a bait, with the parts of colonies closer to the bait showing faster expansion and occupying larger areas with lower cell density compared to the distal portions. Second, the baited expansion phenotype was enhanced when we inoculated the medium with a bait colony and allowed it to grow for 72 hours before inoculating the medium with *Pst* DC3000, which resulted in the proximal portion colonizing a larger area than with simultaneous inoculation. Finally, *Pst* DC3000 induced expansion occurred even after the bait colony was removed from the medium. However, in this case the directional bias was not observed, suggesting that stimulating factor was present, but that it diffuses quickly without the presence of bait cells to maintain the gradient structure. Collectively, these results indicate that the bait cells produce and maintain a soluble gradient of an unknown factor in the medium that is sensed by and induces *Pst* DC3000 directional motility. We did not see any motility change when mixing the supernatant of bait culture into the swimming agar, suggesting the production of such cue/signal might have additional requirements that are only present in solid or semi-solid media.

Bacteria use multi-protein chemotaxis systems for directional control, enabling them to move up or down chemical gradients (Wolfe and Berg 1989; Sampedro et al. 2015). We found little evidence for sub-population-specific regulation of any chemotaxis receptors, except for PSPTO_2448, a methyl-accepting chemotaxis protein that was strongly downregulated in the bait-proximal population (Supplemental Table 1). We attempted to test the role of chemotaxis by genetic manipulation of chemotaxis genes such as *cheA2* and *cheY2* (PSPTO_1980), with the expectation that altering their expression would make the cells unresponsive to the bait. However, deletion of these chemotaxis genes as well as overexpression of *cheY2* led to non-motile phenotype and we could not conclude whether chemotaxis is involved with the baited expansion behavior.

We don’t yet know whether this behavior is conserved among other *Pseudomonas* species or bacteria in general. We used *Pst* DC3000 baited by non-motile *E. coli* as the primary pair in our experiments, but we have no evidence to suggest that *Pst* DC3000 baited expansion is dependent on bait species. We have observed similar baited expansion behaviors when *Pst* DC3000 was baited by *Xanthomonas campestris* (Reeve et al. manuscript submitted) or *Erwinia amylovora* (data not shown), suggesting that *Pst* DC3000 is not responding to a stimulus specific to enteric bacteria (i.e., *E. coli* or *D. dianthicola*). Remarkably, *Pst* DC3000 doesn’t execute baited expansion behavior when presented with a second clone of *Pst* DC3000. This type of selective behavior resembles kin-discrimination behavior, where differential behavior occurs depending on the genetic relationships. Additional research is needed to determine whether social induced (baited) expansion is the default response to the presence of neighboring colonies that occurs except when a kinship signal interrupts the induced expansion.

Molybdenum is an essential trace element and Moco biosynthesis is conserved across all domains of life. In bacteria, Moco is a redox cofactor used in over 50 bacterial redox enzymes that have been mainly studied for their roles in anaerobic respiration and virulence, in addition to Moco biosynthesis itself (Leimkühler 2020; Zhong et al. 2020). To our best knowledge, no bacterial molybdoenzymes have documented interactions with the motility regulation pathways, including c-di-GMP, chemotaxis and flagella synthesis. The only case where Moco has been linked to bacterial motility is in *Burkholderia thailandensis* where disrupting *moaA* results in motility and biofilm defects (Andreae et al. 2014). MoaA is a GTP cyclase necessary for the initial steps of Moco biosynthesis (Leimkühler 2020), so it is tempting to speculate about the possibility of cross-talk between Moco biosynthesis and the c-di-GMP-mediated regulation of motility and biofilms. We found that *Pst* DC3000 mutants lacking *moaA* were no longer capable of baited expansion. To test whether this was a consequence of Moco deficiency or loss of another MoaA function, we tested the effect of *moeA* deletion on baited expansion. The *moeA* gene encodes a protein that catalyzes the last step of Moco assembly (Leimkühler 2020). We found that in the absence of *moeA*, *Pst* DC3000 lost its baited expansion abilities, phenocopying the *moaA* mutant with regard to socially-induced motility, but did not have the reduced baseline motility of the *moaA* mutant. This indicates that Moco is needed for this specific socially-induced motility rather than general swimming motility. Determining how Moco affects socially-induced swimming motility requires further study. In addition, it is a mystery as to whether Moco contributes to the *Pst* DC3000 differential behavior towards kin and non-kin. We suspect that Moco is necessary for some part of a novel motility regulation network that controls baited expansion behavior and not swimming or chemotaxis in general.

We propose that the socially induced motility observed in baited expansion behavior is a form of exploitative competition, in which cells of the baited colony become hyper motile, enabling them to gain access to more territory, nutrients and other resources if uncontested. We only observed baited expansion in interspecies social contexts (Fig. 3), but *Pst* DC3000 did not receive a strong fitness benefit in our assays and the behavior was not species-dependent, which together suggest that cross feeding is unlikely. Additionally, we found no evidence that genes encoding toxin production or T6SS were differentially expressed between subpopulations (Supplement Table S1). However, we cannot rule out the possibility that *Pst* DC3000 might engage in this sort of interference competition after coming into contact with baiting cells.

Possibly one of the most interesting qualities of *Pst* DC3000 baited expansion is that it seems to be a type of delayed gratification game. Under this behavioral strategy, *Pst* DC3000 engages in energetically costly movement without an immediate fitness return, presumably to scramble for resources in an effort to deprive competitors and collect potential fitness rewards later.

## Materials and Methods

### Bacteria strains and growth conditions

For general culturing and preparing strains for experiments, *Pst* DC3000 and *D. dianthicola* strains (Table. 1) were grown at 28°C in King’s B (KB) medium (King et al. 1954) and on KB medium solidified with 1.5% (w/v) agar. *E. coli* strains were grown at 37°C in LB medium and on LB medium solidified with 1.5% (w/v) agar (Bertani 1951). Kanamycin, rifampicin and gentamycin were supplied at 50 μg/ml, 50 μg/ml and 10 μg/ml, respectively.

Plasmid DNAs were isolated using Qiagen Miniprep Kit (Qiagen) from overnight cultured *E. coli* TOP10 cells (Invitrogen) and subsequently used to transform *Pst* DC3000 and mutant derivatives by electroporation (Choi et al. 2006).

Hypomotile strain JW1877 *cheA*::*kan* was used as the *E. coli* bait strain (Baba et al. 2006), and *Pst* DC3000 Δ*cheA2* was used as the kin bait strain (Clarke et al. 2016). A hypomotile strain of *D. dianthicola* ME23 Δ*cheA* MotB.p.Y317X was generated by two-step recombination from pR6KT2GW based SacB-encoding suicide vector (Ried and Collmer 1987; Stice et al. 2020).

### Construction of *Pst* DC3000 mutants and *D. dianthicola* ME23 non-motile mutant

We used standard marker exchange mutagenesis to sequentially produce *Pst* DC3000 strains with the PSPTO_5639, *moaA* (PSPTO_2516) and *moeA* (PSPTO_2353) genes deleted (Kvitko and Collmer 2011). Deletion constructs for each gene were made using synthetic linear DNA fragments from Twist Bioscience that contained between 800 to 1000 bp flanking both ends of each gene joined together with *Eco*RI and *Hin*dIII sites added to the 5’ and 3’ends, respectively (Supplement Table S2). Each deletion construct retained first and last six codons of the original gene to minimize polar effects on downstream genes. The DNA fragments were incorporated into pK18MobSacB by restriction enzyme digestion and ligation with T4-ligase (Thermo Fisher Scientific). Each deletion construct was then purified and confirmed by whole plasmid sequencing (Plasmidsaurus) before being used to transform wild-type *Pst* DC3000 by electroporation (Choi et al. 2006) and selection for kanamycin resistant merodiploids. Sucrose was used to select for recombinants that had subsequently eliminated the *sacB*-containing deletion construct plasmid backbone and confirmed to have lost kanamycin resistance. We confirmed the deletion by sequencing PCR products amplified with primers that anneal to sequences flanking the deleted loci. This process was repeated using the *D. dianthicola* ME23 strain to introduce the *cheA* deletion, except an R6K origin plasmid pR6KT2GW was used instead of pBS46.

Overexpression plasmids were constructed with Gateway cloning. An expression cassette of each target gene was ordered as synthetic linear DNA fragments from Twist Bioscience with *attL* sites added to the 5’ and 3’ends. The DNA fragments were incorporated into pBS46 (Swingle et al. 2008a) with Gateway LR Clonase II enzyme mix (Thermo Fisher Scientific).

After whole plasmid sequencing (Plasmidsaurus) confirmation, these plasmids were then used to transform corresponding *Pst* DC3000 deletion mutant strains by electroporation (Choi et al. 2006). The transformants were then selected by Gentamycin resistance.

### Soft agar swimming assays

MOPS minimal media supplied with 0.2% fructose and 0.3% agar was used for all swimming assays (Neidhardt et al. 1974). 5 ml of swimming assay medium was added to 35 x 10 mm petri dish and air dried in a biosafety cabinet hood for exactly 30 minutes before inoculation. *Pst* DC3000 and bait strains overnight cultures were resuspended and adjusted to OD_600_ = 0.4 or OD_600_ = 0.1, respectively, in MOPS liquid media with 0.2% fructose prior to inoculation. For pairwise interaction swimming assays, a 1 μl inoculum from *Pst* DC3000 strain culture was injected into the agar at the geometrical center of the petri dish, and 1 μl inoculum of bait strain culture was injected 15 mm from the *Pst* DC3000 inoculation site. Plates were then transferred to a sealed moisture-enhanced chamber (RH=100%) and incubated at 25°C.

To prepare conditioned swimming medium with bait removed, a 1 μl bait inoculum was spotted on an approximately 5 mm x 5 mm sterile 0.22 μm pore size nitrocellulose membrane (Millipore) positioned at the usual bait inoculation location. These plates were then incubated as described above for 72 hours before the filter discs were removed along with the bait colony. Then a *Pst* DC3000 strain was inoculated, and the plates were incubated following the standard procedure.

For swimming assays with mix-in bait culture, a LB culture of bait *E. coli* strain was grown until stationary phase, washed and then resuspended in MOPs minimal liquid media with 0.2% fructose. We incubated the resuspended culture for another 96 hours and then filtered twice a 0.2 μm pore size syringe filter to remove bacterial cells. We mixed 100 μl of this supernatant with 4.9 ml melted standard swimming agar while in the control group we mixed in 100 μl of fresh MOPs minimal liquid media with 0.2% fructose.

### RNA extraction and quality control

Samples of *Pst* DC3000 subpopulations were collected from colonies grown on swimming medium using 0.13 x 0.33 inch gel extractors (x-tracta™, Promega) and resuspended in TriZol reagent (Invitrogen). RNA was subsequently extracted from each sample in combination with phase lock gel (Quantabio) following manufacturer instructions. An additional DNAse treatment was performed with Ambion DNase I (Invitrogen) before final elution. Three samples were collected for each combination of strain and subpopulation location. RNA sample quality was confirmed using a Qubit (RNA HS kit; Thermo Fisher) to determine concentration and with a Fragment Analyzer (Agilent) to determine RNA integrity at Cornell Biotechnology Resource Center (BRC).

### RNA-seq library preparation and sequence and analysis

Total RNA samples were treated with a NEBNext rRNA depletion kit (NEB) to remove rRNA. UDI-barcoded RNAseq libraries were generated with the NEBNext Ultra II RNA Library Prep Kit (NEB). Each library was quantified with a Qubit (dsDNA HS kit; Thermo Fisher) and the size distribution was determined with a Fragment Analyzer (Agilent) prior to pooling. Libraries were then sequenced on an Illumina instrument, which produced 10M reads per library.

### RNA-seq analysis

Low quality and adaptor sequences were trimmed with TrimGalore (v0.6.0), a wrapper for cutadpt and fastQC, with the parameters as following: -j 1 -e 0.1 –nextseq-trim=20 -O 1 -a AGATCGGAAGAGC –length 50 –fastqc. Remaining reads were mapped to the *Pst* DC3000 reference genome (Buell et al. 2003) with STAR (v2.7.0e). Normalized counts of reads and statistical analysis of differential gene expression was then carried out with SARTools and DESeq2 (v1.26.0). Subsequent analyses and plotting were performed in R (4.3.2) using package ggplot2 (v.3.4.2), ggbreak (v.0.1.2) and tidyr (v.1.3.0). Principal component analysis (PCA) was carried out independent of DESeq2 using normalized read counts of 1756 non-rRNA genes with at least one significant difference over all pairwise tests (Supplemental Table S1).

### Quantitative Real-Time PCR

We carried out reverse transcription with RNA prepared for transcriptome analysis using an iScript cDNA synthesis kit (Bio-Rad, Hercules, CA). The cDNA product was diluted 10-fold with water and we used 1 μl per technical replicate for quantitative PCR analysis. The qRT-PCR was run with Bio-Rad CFX Connect real-time PCR detection system in combination with SsoAdvanced universal SYBR green (Bio-Rad) following manufacturer-provided instructions. Cycling protocol was set as follows: 95 °C for 3 minutes for initial denaturing, then 40 cycles of 15 seconds at 95 °C, 30 seconds at 52°C and 30 seconds at 60°C. Housekeeping gene *gyrA* was used for baseline standardization (Ferreira et al. 2006; Smith et al. 2018). All primer sequences of *Pst* DC3000 genes are listed in Supplemental table S2.

## Acknowledgements

We thank Drs. Hudson Kern Reeve, Paul Stodghill, Darragh Hare, Adam Bogdanove and Steven Winans for helpful discussions and feedback. We thank Zhongmeng Bao for valuable advice through all cloning processes involved in this work.

Mention of trade names or commercial products in this publication is solely for the purpose of providing specific information and does not imply recommendation or endorsement by the United States Department of Agriculture. USDA is an equal opportunity provider and employer.

## References

Andreae, C. A., Titball, R. W., and Butler, C. S. 2014. Influence of the molybdenum cofactor biosynthesis on anaerobic respiration, biofilm formation and motility in Burkholderia thailandensis. Res Microbiol. 165:41–49

Aoki, S. K., Diner, E. J., de Roodenbeke, C. t’Kint, Burgess, B. R., Poole, S. J., Braaten, B. A., Jones, A. M., Webb, J. S., Hayes, C. S., Cotter, P. A., and Low, D. A. 2010. A widespread family of polymorphic contact-dependent toxin delivery systems in bacteria. Nature. 468:439–442

Aoki, S. K., Pamma, R., Hernday, A. D., Bickham, J. E., Braaten, B. A., and Low, D. A. 2005. Contact-Dependent Inhibition of Growth in Escherichia coli. Science. 309:1245–1248

Baba, T., Ara, T., Hasegawa, M., Takai, Y., Okumura, Y., Baba, M., Datsenko, K. A., Tomita, M., Wanner, B. L., and Mori, H. 2006. Construction of Escherichia coli K-12 in-frame, single-gene knockout mutants: the Keio collection. Mol Syst Biol. 2:2006.0008

Bodman, S. B. von, Willey, J. M., and Diggle, S. P. 2008. Cell-Cell Communication in Bacteria: United We Stand. Journal of Bacteriology. 190:4377–4391

Buell, C. R., Joardar, V., Lindeberg, M., Selengut, J., Paulsen, I. T., Gwinn, M. L., Dodson, R. J., Deboy, R. T., Durkin, A. S., Kolonay, J. F., Madupu, R., Daugherty, S., Brinkac, L., Beanan, M. J., Haft, D. H., Nelson, W. C., Davidsen, T., Zafar, N., Zhou, L., Liu, J., Yuan, Q., Khouri, H., Fedorova, N., Tran, B., Russell, D., Berry, K., Utterback, T., Van Aken, S. E., Feldblyum, T. V., D’Ascenzo, M., Deng, W.-L., Ramos, A. R., Alfano, J. R., Cartinhour, S., Chatterjee, A. K., Delaney, T. P., Lazarowitz, S. G., Martin, G. B., Schneider, D. J., Tang, X., Bender, C. L., White, O., Fraser, C. M., and Collmer, A. 2003. The complete genome sequence of the Arabidopsis and tomato pathogen Pseudomonas syringae pv. tomato DC3000. Proc Natl Acad Sci U S A. 100:10181–10186

Choi, K.-H., Kumar, A., and Schweizer, H. P. 2006. A 10-min method for preparation of highly electrocompetent Pseudomonas aeruginosa cells: application for DNA fragment transfer between chromosomes and plasmid transformation. J Microbiol Methods. 64:391–397

Clarke, C. R., Hayes, B. W., Runde, B. J., Markel, E., Swingle, B. M., and Vinatzer, B. A. 2016. Comparative genomics of Pseudomonas syringae pathovar tomato reveals novel chemotaxis pathways associated with motility and plant pathogenicity. PeerJ. 4

Cornforth, D. M., and Foster, K. R. 2013. Competition sensing: the social side of bacterial stress responses. Nature Reviews Microbiology. 11:nrmicro2977

Garbeva, P., Silby, M. W., Raaijmakers, J. M., Levy, S. B., and Boer, W. D. 2011. Transcriptional and antagonistic responses of Pseudomonas fluorescens Pf0-1 to phylogenetically different bacterial competitors. The ISME Journal; London. 5:973–85

Gibbs, K. A., Urbanowski, M. L., and Greenberg, E. P. 2008. Genetic Determinants of Self Identity and Social Recognition in Bacteria. Science. 321:256–259

Haapalainen, M., Mosorin, H., Dorati, F., Wu, R.-F., Roine, E., Taira, S., Nissinen, R., Mattinen, L., Jackson, R., Pirhonen, M., and Lin, N.-C. 2012. Hcp2, a Secreted Protein of the Phytopathogen Pseudomonas syringae pv. Tomato DC3000, Is Required for Fitness for Competition against Bacteria and Yeasts. J Bacteriol. 194:4810–4822

Hayes, C. S., Koskiniemi, S., Ruhe, Z. C., Poole, S. J., and Low, D. A. 2014. Mechanisms and Biological Roles of Contact-Dependent Growth Inhibition Systems. Cold Spring Harb Perspect Med. 4

Hosni, T., Moretti, C., Devescovi, G., Suarez-Moreno, Z. R., Fatmi, M. B., Guarnaccia, C., Pongor, S., Onofri, A., Buonaurio, R., and Venturi, V. 2011. Sharing of quorum-sensing signals and role of interspecies communities in a bacterial plant disease. ISME J. 5:1857–1870

Humphries, J., Xiong, L., Liu, J., Prindle, A., Yuan, F., Arjes, H. A., Tsimring, L., and Süel, G. M. 2017. Species-Independent Attraction to Biofilms through Electrical Signaling. Cell. 168:200–209.e12

Jones, S. E., Ho, L., Rees, C. A., Hill, J. E., Nodwell, J. R., and Elliot, M. A. 2017. Streptomyces exploration is triggered by fungal interactions and volatile signals T. Mignot, ed. eLife. 6:e21738

Kvitko, B. H., and Collmer, A. 2011. Construction of Pseudomonas syringae pv. tomato DC3000 mutant and polymutant strains. Methods Mol Biol. 712:109–128

Leimkühler, S. 2020. The biosynthesis of the molybdenum cofactors in Escherichia coli. Environmental Microbiology. 22:2007–2026

Limoli, D. H., Warren, E. A., Yarrington, K. D., Donegan, N. P., Cheung, A. L., and O’Toole, G. 2019. Interspecies interactions induce exploratory motility in Pseudomonas aeruginosa D.K. Newman, ed. eLife. 8:e47365

Liu, J., Martinez-Corral, R., Prindle, A., Lee, D. D., Larkin, J., Gabalda-Sagarra, M., Garcia-Ojalvo, J., and Süel, G. M. 2017. Coupling between distant biofilms and emergence of nutrient time-sharing. Science. :eaah4204

Liu, Y., Kyle, S., and Straight, P. D. 2018. Antibiotic Stimulation of a Bacillus subtilis Migratory Response. mSphere. 3

Ma, X., Perna, N. T., Glasner, J. D., Hao, J., Johnson, S., Nasaruddin, A. S., Charkowski, A. O., Wu, S., Fei, Z., Perry, K. L., Stodghill, P., and Swingle, B. 2019. Complete Genome Sequence of Dickeya dianthicola ME23, a Pathogen Causing Blackleg and Soft Rot Diseases of Potato. Microbiol Resour Announc. 8:e01526–18

Morris, C. E., Sands, D. C., Vinatzer, B. A., Glaux, C., Guilbaud, C., Buffière, A., Yan, S., Dominguez, H., and Thompson, B. M. 2008. The life history of the plant pathogen Pseudomonas syringae is linked to the water cycle. ISME J. 2:321–334

Neidhardt, F. C., Bloch, P. L., and Smith, D. F. 1974. Culture Medium for Enterobacteria. J Bacteriol. 119:736–747

Reeve, H. K., Hare, D., Yang, Z., Dell’Aquila, M., Liu, Y., and Swingle, B. 2018. Kin-selected avoidance of competition between genetically identical bacteria.

Ried, J. L., and Collmer, A. 1987. An nptI-sacB-sacR cartridge for constructing directed, unmarked mutations in gram-negative bacteria by marker exchange-eviction mutagenesis. Gene. 57:239– 246

Sampedro, I., Parales, R. E., Krell, T., and Hill, J. E. 2015. Pseudomonas chemotaxis. FEMS Microbiol Rev. 39:17–46

Schäfer, A., Tauch, A., Jäger, W., Kalinowski, J., Thierbach, G., and Pühler, A. 1994. Small mobilizable multi-purpose cloning vectors derived from the Escherichia coli plasmids pK18 and pK19: selection of defined deletions in the chromosome of Corynebacterium glutamicum. Gene. 145:69–73

Senior, B. W. 1977. The Dienes phenomenon: identification of the determinants of compatibility. J Gen Microbiol. 102:235–244

Shepherdson, E. M. F., and Elliot, M. A. 2022. Cryptic specialized metabolites drive Streptomyces exploration and provide a competitive advantage during growth with other microbes. Proc Natl Acad Sci U S A. 119:e2211052119

Stefanic, P., Kraigher, B., Lyons, N. A., Kolter, R., and Mandic-Mulec, I. 2015. Kin discrimination between sympatric Bacillus subtilis isolates. PNAS. 112:14042–14047

Stice, S. P., Thao, K. K., Khang, C. H., Baltrus, D. A., Dutta, B., and Kvitko, B. H. 2020. Thiosulfinate Tolerance Is a Virulence Strategy of an Atypical Bacterial Pathogen of Onion. Curr Biol. 30:3130–3140.e6

Swingle, B., Thete, D., Moll, M., Myers, C. R., Schneider, D. J., and Cartinhour, S. 2008a. Characterization of the PvdS-regulated promoter motif in Pseudomonas syringae pv. tomato DC3000 reveals regulon members and insights regarding PvdS function in other pseudomonads. Mol. Microbiol. 68:871–889

Swingle, B., Thete, D., Moll, M., Myers, C. R., Schneider, D. J., and Cartinhour, S. 2008b. Characterization of the PvdS-regulated promoter motif in Pseudomonas syringae pv. tomato DC3000 reveals regulon members and insights regarding PvdS function in other pseudomonads. Mol. Microbiol. 68:871–889

Waters, C. M., and Bassler, B. L. 2005. Quorum sensing: cell-to-cell communication in bacteria. Annu Rev Cell Dev Biol. 21:319–346

Wolfe, A. J., and Berg, H. C. 1989. Migration of bacteria in semisolid agar. PNAS. 86:6973–6977

Yarrington, K. D., Shendruk, T. N., and Limoli, D. H. 2024. The type IV pilus chemoreceptor PilJ controls chemotaxis of one bacterial species towards another. PLoS Biol. 22:e3002488

Zhong, Q., Kobe, B., and Kappler, U. 2020. Molybdenum Enzymes and How They Support Virulence in Pathogenic Bacteria. Frontiers in Microbiology. 11

